# Evaluation of Nanopore sequencing technology to differentiate *Salmonella* serotypes and serotype variants with the same or closely related antigenic formulae

**DOI:** 10.1101/2020.09.06.274746

**Authors:** Feng Xu, Chongtao Ge, Shaoting Li, Silin Tang, Xingwen Wu, Hao Luo, Xiangyu Deng, Guangtao Zhang, Abigail Stevenson, Robert C. Baker

## Abstract

Our previous study demonstrated that whole genome sequencing (WGS) data generated by Oxford Nanopore Technologies (ONT) can be used for rapid and accurate prediction of *Salmnonella* serotypes. However, one limitation is that established methods for WGS-based serotype prediction cannot differentiate certain serotypes and serotype variants with the same or closely related antigenic formulae. This study aimed to evaluate Nanopore sequencing and corresponding data analysis for differentiation of these serotypes and serotype variants, thus overcoming this limitation. Five workflows that combined different flow cells, library construction methods and basecaller models were evaluated and compared. The workflow that consisted of the R9 flow cell, rapid sequencing library construction kit and guppy basecaller with base modified model performed best for Single Nucleotide Polymorphism (SNP) analysis. With this workflow, as high as 99.98% matched the identity of the assembled genomes and only less than five high quality SNPs (hqSNPs) between ONT and Illumina sequencing data were achieved. SNP typing allowed differentiation of Choleraesuis*sensu stricto*, Choleraesuis var. Kunzendorf, Choleraesuis var. Decatur, Paratyphi C, and Typhisuis that share the same antigenic formula 6,7:c:1,5. Prophage prediction further distinguished Orion var. 15^+^ and Orion var. 15^+^, 34^+^. Our study improves the readiness of ONT as a *Salmonella* subtyping and source tracking tool for food industry applications.

**Highlights:**

- *Salmonella* serotypes or serotype variants with the same antigenic formula were differentiated by SNP typing.
- Nanopore sequencing followed by phage prediction identified the *Salmonella* serotype variants caused by phage conversion.
- The latest ONT technology is capable of high fidelity SNP typing of *Salmonella*.

## 1. Introduction

*Salmonella* remains an important concern in the food industry since this foodborne pathogen continues to cause global public health and economic impact. Thus, it is imperative to reinforce *Salmonella* control measures in the food industry (GMA, 2009). Rapid methods for subtyping strains beyond the species level to serotype are essential to facilitate contamination incident investigations (Olaimat and Holley, 2012; Shi et al., 2015). There are more than 2,600 *Salmonella* serotypes currently described in the White-Kauffmann-Le Minor scheme (Grimont and Weill, 2007; Dieckmann and Malorny, 2011). Whole Genome Sequencing (WGS) based tools have now emerged as an alternative method for *Salmonella* serotyping (Zhang et al., 2015; Yoshida et al., 2016; Yachison et al., 2017; Ibrahim and Morin, 2018; Diep et al., 2019; Tang et al., 2019; Zhang et al., 2019; Uelze et al., 2020) and have been suggested to be the new gold standard (Banerji et al., 2020).

Oxford Nanopore Technologies (ONT) sequencing generates long-read sequences with rapid turnaround and has the potential for rapid identification of food-borne pathogens during routine monitoring and incident investigations in the food industry. Challenges in the basecalling accuracy of nanopore sequencing have been identified compared with short-read sequencing technologies (Amarasinghe et al., 2020). Errors caused by signal interpretation during basecalling are dominated by insertions and deletions (indels), especially in homopolymeric regions (Gargis et al, 2019). An average of 95% accuracy of raw read (Jain et al., 2017, 2018) and 1.06 errors per Kbp assembly (Lang et al., 2020) were recently reported.

*In silico* serotype prediction using ONT sequences was systematically evaluated in our previous study, and all the tested *Salmonella* isolates representing 34 serotypes were correctly predicted at the level of antigentic profile (Xu et al., 2020). Comparison with the conventional serotyping method using antisera combined with biochemical tests, identified a limitation that the established WGS methods, with either Illumina or ONT data, were unable to differentiate serotypes that share the same or related antigenic formulae.

Serotypes with antigenic formula 6,7:c:1,5 including Paratyphi C (Vi + or Vi–), Typhisuis, Choleraesuis *sensu stricto*, Choleraesuis var. Kunzendorf and Choleraesuis var. Decatur, require additional biochemical tests to differentiate between them in conventional serotyping (Grimont and Weill, 2007). Choleraesuis is a serotype adapted to swine. This serotype has caused serious disease outbreaks in swine, including two in Denmark in 1999 – 2000 and 2012 – 2013 (Leekitcharoenphon et al., 2019). It also has a propensity to cause extraintestinal infections in humans (Chiu et al., 2004; Sirichote et al., 2010). Choleraesuis var. Kunzendorf is responsible for the majority of outbreaks among swine (Leekitcharoenphon et al., 2019). Serotype Orion has two closely related variants, Orion var. 15^+^ and Orion var. 15^+^, 34^+^. Phage conversion by □_15_ or □_34_ is needed to differentiate these two variants (Grimont and Weill, 2007). Orion has been isolated from a variety of hosts worldwide, including humans (Cabrera et al., 2006; Trafny et al., 2006), cattle (Alam et al., 2009), ducks (Rampersad et al., 2008), dogs (Gupta et al., 2016) and birds (Münch et al., 2012). Orion var.15^+^ and Orion var. 15^+^, 34^+^ are more likely to be associated with poultry (Corry et al., 2002; McWhorter et al., 2015). Additional subtyping analysis is needed to differentiate these variants.

When the genetic determinant for a particular biotype or variant is known, it makes an effective target for futher differentiation. For example, SeqSero2 uses a Single Nucleotide Polymorphism (SNP) that inactivates tartrate fermentation to differentiate the *S*. Paratyphi B pathogen and a 7-bp deletion to identify O5-variants of *S*. Typhimurium (Zhang et al., 2019). When such genetic determinants are not available, phylogenetic analysis, such as SNP typing and core/whole genome Multilocus Sequence Typing (MLST), can be used to differentiate biotypes and serotype. Nair et al. grouped the 6,7:c:1,5 biotypes of *S. enterica* subspecies *enterica* including Paratyphi C, Typhisuis, Choleraesuis *sensu stricto*, Cholerasuis var. Kunzendorf into four phyloclusters using the core SNP phylogeny (Nair et al., 2020). The authors also found that Choleraesuis var. Decatur genomes showed more SNP variation between them than was found across all of the remaining biotypes (Nair et al., 2020). With continuous improvement of the sequencing and basecalling accuracy of ONT, application of ONT sequences in SNP analysis has become more applicable (Greig et al., 2019; Taylor et al., 2019).

The goal of this study was to evaluate the combined use of latest improved ONT sequencing and modified subtyping analyses with primary serotype prediction in differentiating representative *Salmonella* serotypes or serotype variants that shared the same or related antigenic formulae.

## 2. Materials and methods

### 2.1. Bacterial strains

Two strains, *S. enterica* Choleraesuis var. Kunzendorf and *S. enterica* Orion var. 15^+^, 34^+^, were obtained from Dr. Martin Weidmann’s laboratory (Cornell University, USA). Details of these strains can be found at www.foodmicrobetracker.com under the isolate IDs FSL R9-0095 and FSL R8-3858.

### 2.2. Genomic DNA extraction and quality control

Strains were incubated for 20 hrs on Trypticase Soy Agar (TSA) at 37°C. Genomic DNA was extracted using a QIAamp DNA mini kit (Qiagen, Hilden, Germany), following the instructions provided by the manufacture. The double-stranded DNA (dsDNA) was quantified using a Qubit 3.0 fluorimeter (Life Technologies, Paisley, UK). The quality of DNA was assessed using a NanoDrop 1000 instrument (Thermo Fisher Scientific, Delaware, USA).

### 2.3. Illumina sequencing and genome assembly

Illumina sequences were performed at the Beijing Genomics Institute (BGI, Shenzhen, China). The library for sequencing was constructed by BGI and sequenced using a HiSeq X Ten platform (Illumina, San Diego, CA, USA) to 2 × 150 cycles. Genomes were assembled by the SOAPdenovo v1.05 (URL: http://soap.genomics.org.cn).

### 2.4. Oxford Nanopore sequencing and genome assembly

Five workflows with combinations of different flow cells, library construction methods and basecaller models were used in this study for comparison purposes (Supplementary Table 1). The DNA library was prepared with library construction kits following the manufacturer’s instructions as indicated in each workflow. For flow cell, compared with commercial available R9 (version 9.4.1), R10 (version 10.0, early access) is a new design of nanopore, with a longer barrel and dual reader head. Libraries were sequenced with qualified R9 and R10 flow cells (active pores number >= 800) on a GridION sequencer (Oxford Nanopore Technologies, Oxford, UK). Real-time basecalling was performed using Guppy version 3.2.6 with the corresponding basecalling model as indicated in each workflow, which was integrated in the MinKNOW software v3.5.4. Fastq data were obtained for further analysis. Adaptor sequences were trimmed with Porechop v0.2.3 (URL: https://github.com/rrwick/Porechop), and the quality of trimmed data was assessed using NanoStat v1.1.2 (De Coster et al., 2018). Filtlong v0.2.0 (URL: https://github.com/rrwick/Filtlong) was applied to remove reads shorter than 1,000 bp. Genomes were *de novo* assembled by Wtdbg2 v2.4 (Ruan and Li, 2019) with default parameters. The assembled genomes were corrected using Racon v1.3.3 (Vaser et al. 2017). Consensus was obtained from one round of Medaka version 0.8.0 (https://github.com/nanoporetech/medaka). QUAST v5.0.2 was applied to assess the assembled contigs (Gurevich et al., 2013). DNAdiff v1.3 (MUMmer version, Kurtz et al., 2004) was used to evaluate the base level comparison between ONT and Illumina assemblies.

**Table 1.**
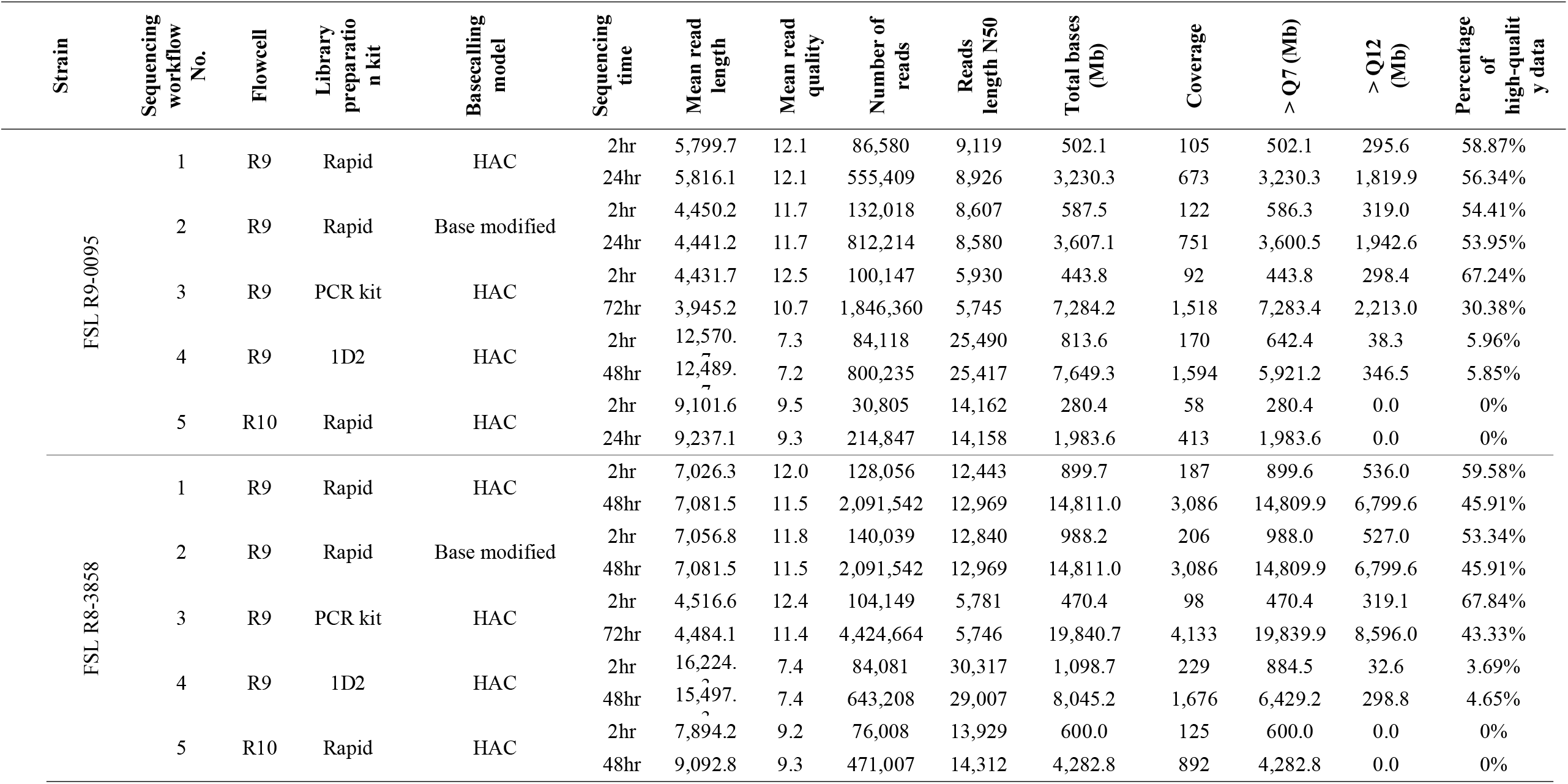
Overview of ONT raw reads generated using different sequencing workflows.

### 2.5. Phylogenetic analysis

#### 2.5.1. kSNP tree

A k-mer based approach, kSNP3 (Gardner et al., 2015) was used to assess the clustering of isolates sharing the same antigenic formula with FSL R9-0095 and isolates sharing closely related antigenic formulae with FSL R8-3858. Kchooser was used to determine an optimum k-mer size (Carroll et al., 2017). A parsimony tree was generated by kSNP3 using the set of core SNPs identified. Clade robustness was assessed using a bootstrap analysis with 1,000 replicates (Felsenstein, 1985). Twenty-six draft genome sequences were downloaded from GenBank and further confirmed by SeqSero2 (Zhang et al., 2019) and SISTR (Yoshida et al., 2016) for the phylogenetic analysis of strain FSL R9-0095 (Supplementary Table 2). These draft genome sequences included six strains of Typhisuis, eight strains of Paratyphi C, two strains of Choleraesuis *sensu stricto*, three strains of Choleraesuis var. Decatur, and seven strains of Choeraesuis var. Kunzendorf. The Illumina sequences of strain FSL R9-0095 and the corresponding ONT sequences generated using all the sequencing workflows were analyzed through kSNP analysis. Twenty-seven draft genomes were downloaded from GenBank and further confirmed by SeqSero2 and SISTR for the phylogenetic analysis of strain FSL R8-3858 (Supplementary Table 2). These draft genomes included 16 strains of Orion, seven strains of Orion var. 15^+^, and four strains of Orion var. 15^+^, 34^+^. The Illumina sequences of strain FSL R8-3858 and the corresponding ONT sequences generated using all the sequencing workflows were analyzed through kSNP analysis.

**Table 2.** Overview of ONT assembled contigs through data analysis pipeline 1.

| Strain | Sequencing workflow No. | Flowcell | Library Preparation kit | Basecalling model | Sequencing time | Total length (bp) | Contigs Numbers | N50 Length (bp) | SNPs | InDels | Matching identity (%) | High quality SNPs |
| --- | --- | --- | --- | --- | --- | --- | --- | --- | --- | --- | --- | --- |
| FSL R9-0095 | 1 | R9 | Rapid | HAC | 2hr | 4,792,171 | 3 | 4,736,332 | 4,778 | 3,116 | 99.83 | 4 |
|  |  |  |  |  | 24hr | 4,789,608 | 2 | 4,741,208 | 4,492 | 2,534 | 99.85 | 2 |
|  | 2 | R9 | Rapid | Base modified | 2hr | 4,825,067 | 4 | 4,740,589 | 31 | 901 | 99.98 | 0 |
|  |  |  |  |  | 24hr | 4,741,424 | 1 | 4,741,424 | 29 | 652 | 99.98 | 0 |
|  | 3 | R9 | PCR kit | HAC | 2hr | 4,667,484 | 49 | 183,328 | 42 | 493 | 99.99 | 0 |
|  |  |  |  |  | 72hr | 4,561,879 | 48 | 174,019 | 245 | 1,001 | 99.97 | 0 |
|  | 4 | R9 | 1D2 | HAC | 2hr | 4,742,094 | 1 | 4,742,094 | 5,043 | 5,840 | 99.77 | 1 |
|  |  |  |  |  | 48hr | 4,790,286 | 2 | 4,741,889 | 5,200 | 4,911 | 99.78 | 4 |
|  | 5 | R10 | Rapid | HAC | 2hr | 4,790,793 | 2 | 4,742,687 | 619 | 1,672 | 99.95 | 1 |
|  |  |  |  |  | 24hr | 4,783,849 | 2 | 4,735,698 | 811 | 2,179 | 99.93 | 1 |
| FSL R8-3858 | 1 | R9 | Rapid | HAC | 2hr | 4,665,249 | 1 | 4,665,249 | 4,831 | 2,617 | 99.84 | 100 |
|  |  |  |  |  | 48hr | 4,665,134 | 1 | 4,665,134 | 4,854 | 2,705 | 99.83 | 73 |
|  | 2 | R9 | Rapid | Base modified | 2hr | 4,666,028 | 1 | 4,666,028 | 33 | 677 | 99.98 | 1 |
|  |  |  |  |  | 48hr | 4,666,136 | 1 | 4,666,136 | 38 | 666 | 99.98 | 1 |
|  | 3 | R9 | PCR kit | HAC | 2hr | 4,583,190 | 31 | 273,714 | 56 | 483 | 99.98 | 1 |
|  |  |  |  |  | 72hr | 4,500,549 | 33 | 180,233 | 59 | 407 | 99.99 | 1 |
|  | 4 | R9 | 1D2 | HAC | 2hr | 4,666,639 | 1 | 4,666,639 | 5,605 | 6,524 | 99.74 | 69 |
|  |  |  |  |  | 48hr | 4,698,102 | 2 | 4,665,652 | 5,964 | 6,724 | 99.72 | 61 |
|  | 5 | R10 | Rapid | HAC | 2hr | 4,666,923 | 1 | 4,666,923 | 691 | 1,919 | 99.94 | 4 |
|  |  |  |  |  | 48hr | 4,651,530 | 2 | 3,508,306 | 608 | 1,873 | 99.95 | 3 |

#### 2.5.2. CFSAN high quality SNP (hqSNP) pipeline

The hqSNP pipeline developed by the Center for Food Safety and Applied Nutrition (CFSAN SNP Pipeline v.1.0.0/FDA) was used for SNP calling of *Salmonella* strains with default quality filters (Davis et al., 2015). Read mapping was performed using Burrows-Wheeler Aligner (v0.7.17-r1188) (BWA-MEM) with the -x ont2d option (Li and Durbin, 2010; Hyeon et al., 2018) for the ONT sequences. An SNP matrix and alignment concatenated SNP were produced with customized Python scripts.

#### 2.5.3. Phylogenetic tree generated from CFSAN hqSNP analysis

Maximum likelihood (ML) phylogenetic trees were built using PhyML v3.3 based on the hqSNP matrices and visualized with Figtree v1.4.4 (http://tree.bio.ed.ac.uk/software/figtree/). All the phylogenetic trees shown in this paper were midpoint rooted.

### 2.6. Phage detection

All the Illumina data of Orion and its varirants, as well as the assembled genomes of strain FSL R8-3858 sequenced using either Illumina or ONT, were submitted to the Phage Search Tool – Enhanced Release (PHASTER API website, http://phaster.ca). Results were extracted from the files returned from the server (Zhou et al., 2011; Arndt et al., 2016; Worley et al., 2018). Sequences classified by PHASTER as questionable (score between 70 and 90), intact (score > 90) or incomplete (score < 70) were all recorded.

## 3. Results and discussion

### 3.1. Overview of ONT raw reads

The same ONT fast5 data were used for basecalling by the high accuracy (HAC) model and base modified model (workflow 1 vs workflow 2) (Supplementaryable 1). The data with quality score >= 12 were considered high quality data. The mean read length, mean read quality and percentage of high-quality data (high quality data/total bases) were similar between data generated by these two basecalling models (Table 1). The read length of sequences generated from workflow 3 was close to 4 Kbp (Table 1). One strand of dsDNA prepared by the 1D2 library preparation kit was sequenced followed by its complementary strand for workflow 4. This workflow generated sequences with the longest mean read length (Table 1). However, its mean read quality of the sequences was the lowest (Table 1).

Two equal aliquots of the same batch of extracted genomic DNA were sequenced with the R9 and R10 flow cells for workflows 1 and 5, respectively. The yield of the R9 flow cell was higher than that of the R10 flow cell (Table 1). The mean read length of sequences from the R9 flow cell were shorter than that from the R10 flow cell (Table 1). The mean read quality of sequences from the R9 flow cell were higher than that from the R10 flow cell (Table 1). The percentage of high quality data of sequences from the R9 flow cell were higher than that from the R10 flow cell (Table 1).

According to the previous study sequences produced in two hrs from ONT were sufficient for *Salmonella* serotype prediction (Xu et al., 2020). Therefore, the raw ONT sequences from the first two hrs of sequencing time were extracted to investigate whether they were sufficient for additional subtyping analysis. About 100× genome coverage was achieved after two hrs sequencing, except with workflow 5 which used the R10 flow cell for FSL R9-0095 sequencing (58×, Table 1). This might have resulted from a dramatic decrease of active pores on the R10 from 1,223 to 155 after the library was loaded, although it increased after 1.5 hrs. Overall, comparisons of the sequencing data yield, mean read length, mean quality score and high-quality data percentage from different workflow using two hrs sequencing data were in line with those using the data of full runs (24, 48, 72 hrs, Table 1).

### 3.2. Overview of ONT assembled contigs

The ONT sequencing data were assembled using Wtdbg2, and corrected by 2 rounds of Racon. Consensus were then obtained by one round of Medaka (Supplementary Figure 1). The overview of the assembled contigs through the reads obtained on GridION are summarized below (Table 2). The contigs generated from Illumina sequencing were used as a benchmark for comparisons. In general, higher matching identity (> 99.90%) and fewer SNPs by DNAdiff (< 1,000) were obtained from workflows 2, 3 and 5 compared with workflows 1 or 4 (< 99.90% of matching identity and >1,000 of SNPs by DNAdiff). DNA libraries prepared with the rapid kit (workflows 1, 2 and 5) and 1D2 kit (workflow 4) generated fewer contigs (< 5) and higher N50 length (> 4.6 Mbp) in general, except for the 48 hrs data obtained with workflow 5 for FSL R8-3858 sequencing (N50 = 3.51 Mbp). The DNA library prepared using the PCR kit (workflow 3) generated more contigs and shorter N50 length (Table 2). The reason could be fragmentation of genomic DNA followed by a PCR amplification step during the library preparation. Taylor et al. (2019) reported that four hrs ONT sequences generated full-length genomes with an average identity of 99.87% for *S. Bareilly* and 99.89% for *Escherichia coli* in comparison to the respective Illumina references. With the accuracy improvement of ONT sequencing, especially for the basecaller and flow cell, the highest matching identity was improved to 99.99% as demonstrated using sequencing workflow 3 with the PCR kit in this study (Table 2).

**Figure 1.**
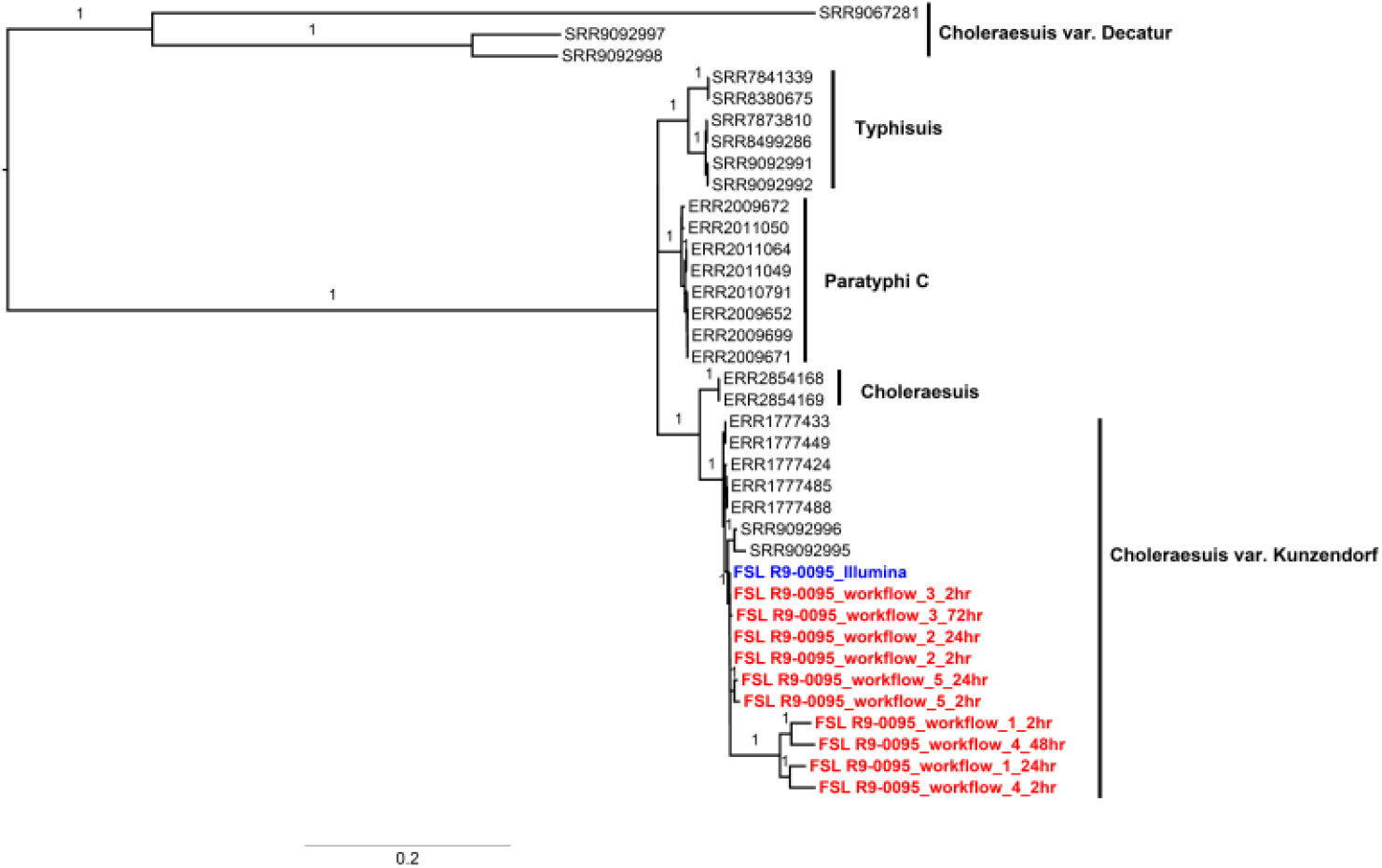
Maximum parsimony tree of the serotype formula 6,7:c:1,5 based on k-mer-based SNP analysis. The tree was built using kSNP3 with the core SNPs identified among both Illumina and ONT sequences of the strain FSL R9-0095 and 26 sequences downloaded from the NCBI. Clades of Typhisuis, Paratyphi C, Choleraesuis, Choleraesuis var. Kunzendorf, Choleraesuis var. Decatur were annotated separately. Only high support in the analysis (bootstrap >= 90%) are labeled in this tree. The tree is midpoint rooted and the scale axis is provided below the tree.

Two sequencing workflows were tested to reduce the impact of methylation, as an average of 95% of the discrepant positions between the sequencing technologies (ONT *versus* Illumina) were caused by methylations (Greig et al., 2019). These were workflow 2 (R9 flow cell + rapid kit + base modified model basecaller) and workflow 3 (R9 flow cell + PCR kit + HAC model basecaller). The base modified model basecaller was trained with human and *E. coli* reads, which allowed for calling 5-methylcytosine (5mC) and N6-methyladenine (6mA) modified bases. In comparison with workflow 1, the total SNP loci generated from workflow 2 decreased from 4492 – 4854 to 29 – 38 for both tested strains (Table 2). Since workflow 3 included a PCR amplification step in library preparation, it could exclude methylations from the library (Liu et al., 2019). Compared with workflow 1, the total SNP loci decreased to 42 – 245 for both tested strains (Table 2). Workflow 4 generated similar SNP numbers to that of workflow 1, ranging from 4,400 to 6,000, which suggested that sequencing both DNA strands might not improve accuracy (Table 2). Despite the low data quantity and genome coverage, fewer SNPs were obtained by sequencing on R10 flow cells (workflow 5) compared to R9 flow cells (workflow 1). R10 flow cell had a longer barrel and dual reader head, enabling improved resolution of homopolymeric regions and improving the consensus accuracy of ONT sequencing data (https://nanoporetech.com/about-us/news/r103-newest-nanopore-high-accuracy-nanopore-sequencing-now-available-store). Current results indicated that flow cell R10 flow cell did improve sequencing accuracy.

### 3.3. Differentialtion of serotypes and serotype variants that share the same antigenic formula

#### 3.3.1 SNP analysis

Isolates belonging to each of the three serotypes (Typhisuis, Paratyphi C and Choleraesuis) and two serotype variants (Chloeraesuis var. Decatur abd Chloeraesuis var. Kunzendorf) sharing the same antigenic formula 6,7:c:1,5 formed a distinct clade in the kSNP phylogeny with 100% bootstrap support for each clade (Figure 1). This suggested that kSNP analysis using assembled genomes could further differentiate serotypes and variants with this antigenic formula. Genomes of strain FSL R9-0095 sequenced with either the Illumina or ONT platforms fell into the Choleraesuis var. Kunzenforf clade (Figure 1), which indicated that ONT sequencing was equivalent to Illumina for kSNP analysis in identifying these serotypes and serotype variants. Since the SNP numbers generated through sequencing workflows 1 and 4 were higher than those through the other three workflows, the branches of these four corresponding genomes were longer than the other ONT genomes with 100% of bootstrap support (Figure 1). A shorter branch was observed in the subclade through use of workflow 5 (sequencing on the R10 flow cell).

Genomes of strain FSL R9-0095 sequenced on the Illumina platform were used as a reference for read mapping in the analysis of CFSAN hqSNP. ONT sequences generated using the five different workflows and from different sequencing time were analyzed using the CFSAN hqSNP pipeline. The numbers of hqSNP are listed below (Table 2). Sequencing workflows 2 and 3 generated 0 hqSNP between Illumina data and corresponding ONT data for strain FSL R9-0095. Workflow 5 generated 1 hqSNP after two or 24 hrs of sequencing of the same strain. Overall, the 5 sequencing workflows yielded fewer than 10 hqSNPs compared with the corrsponding Illumina sequences (Table 2).

The phylogenetic trees generated by hqSNP using ONT and/or Illumina sequences for strain FSL R9-0095 are presented below (Figure 2a, 2b and 2c). Three phylogenetic trees were constructed. In order to compare ONT and Illumina sequences, the first tree was built using both ONT and Illumina sequences, including Illumina sequences from GenBank, Illumina sequence of the tested isolate, and ONT sequences generated by different workflows and different sequencing time (Figure 2a). In the second tree, one ONT-sequenced isolate was included to replace the corresponding Illumina-sequenced genome to assess hqSNP-based phylogenetic analysis using ONT sequencing (Figure 2b). The third tree was built using Illumina sequences of these isolates only. Strain FSL R9-0095 was correctly placed in the Cholerasuis var. Kunzendorf clade (Figures 2a, 2b and 2c). All the different serotypes or serotype variants under antigenic formula 6,7:c:1,5 were clearly separated into their respective clades, and all the clades were supported by 100% bootstrap. This indicated that the CFSAN hqSNP-based phylogenetic analysis with either ONT or Illumina data was able to differentiate Choleraesuis var. Kunzendorf isolates. In addition, less than 30 mins was needed for library preparation by the rapid library preparation kit, while about three and five hrs were required using the 1D2 and PCR kits, respectively. In summary, workflow 2 (which used the rapid library preparation kit) was the more optimal procedure to deliver the desired phylogenetic analysis in terms of accuracy and tunaround time. The phylogenetic trees constructed from data sets of two sequencing times (two hrs and 24 hrs) using workflow 2 showed identical topology (data not shown), suggesting that ONT sequences collected within two hrs were sufficient for CFSAN hqSNP-based phylogenetic analysis to generate correct variant prediction.

**Figure 2.**
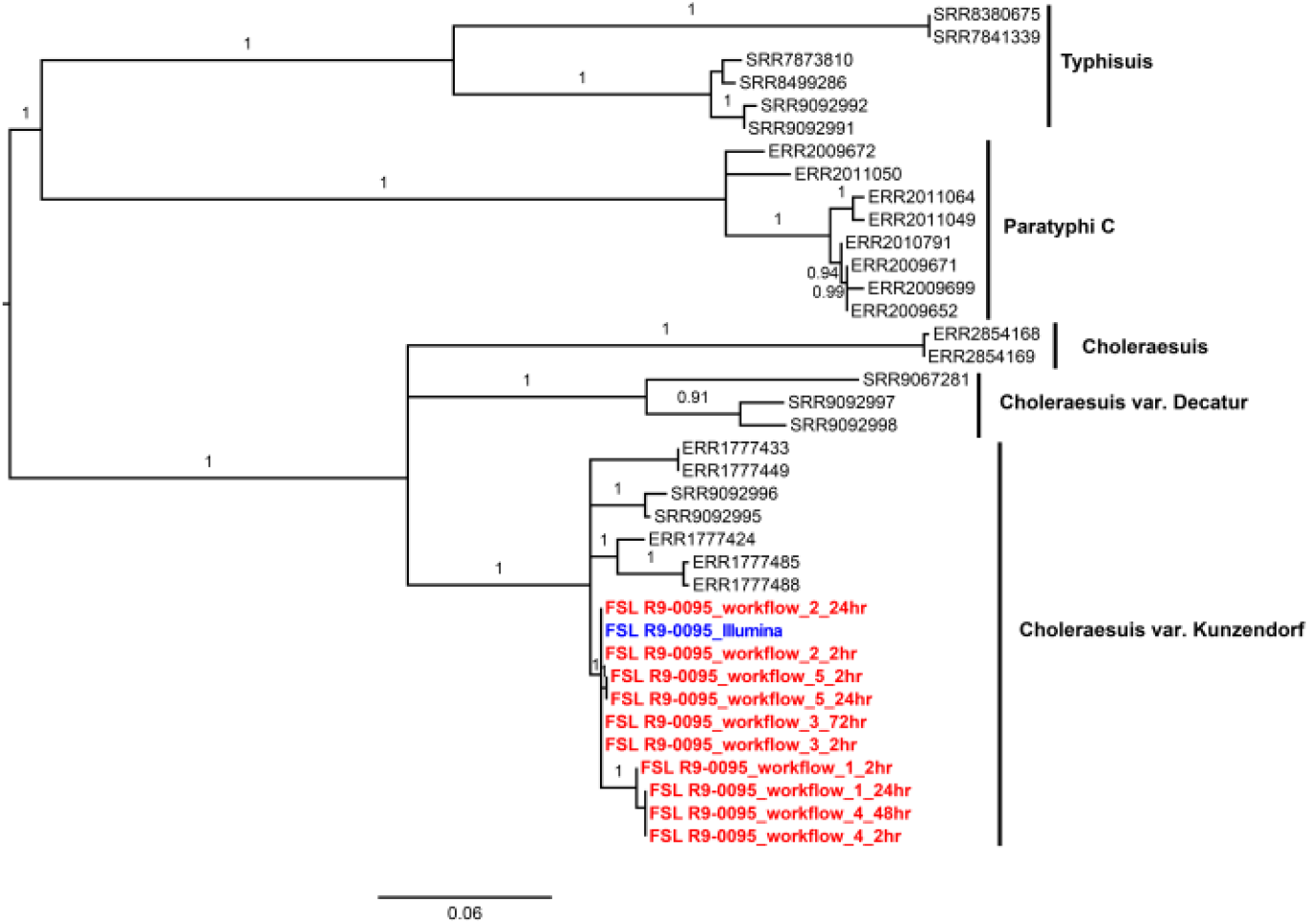
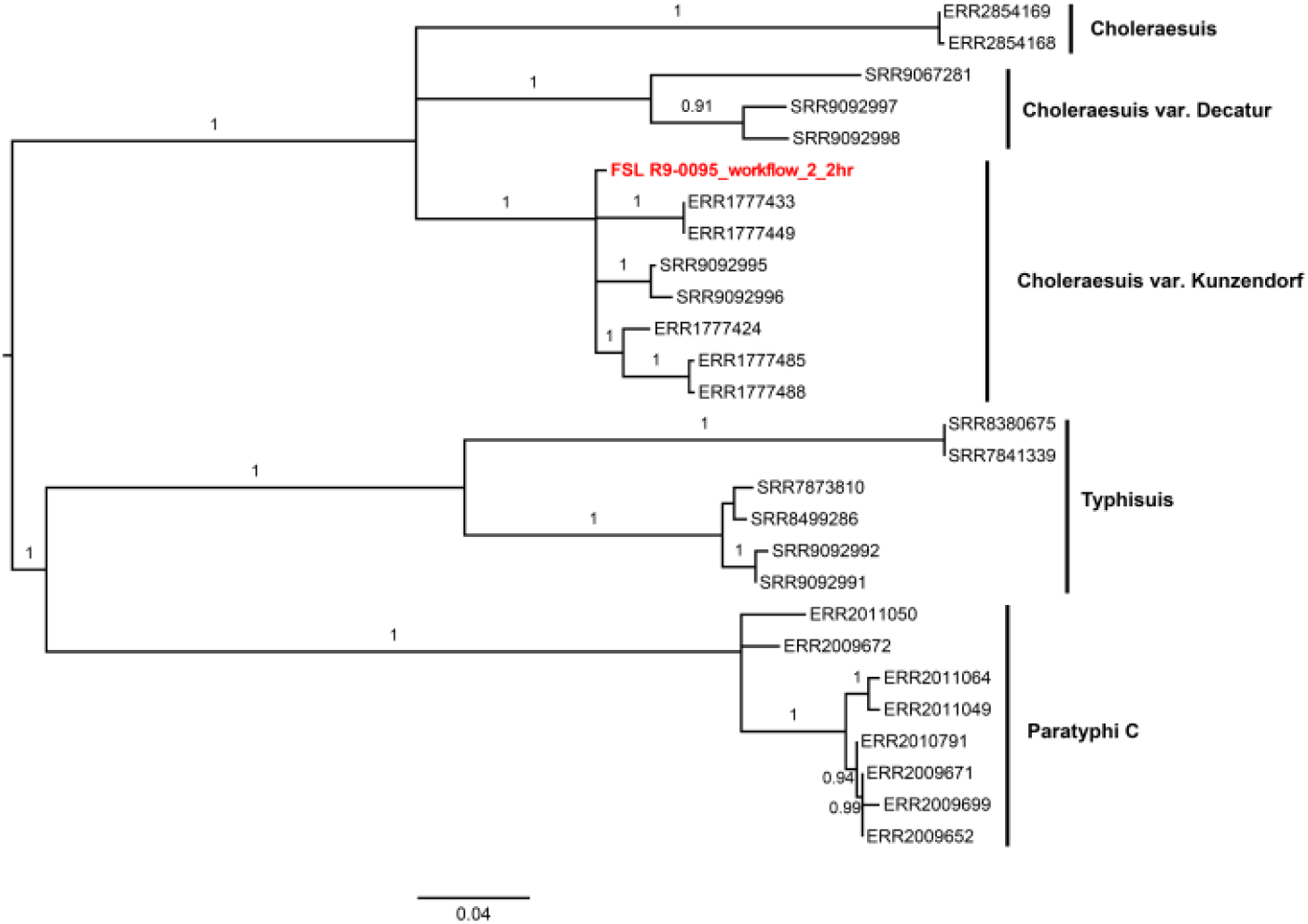
Maximum likelihood phylogenetic tree of the serotype formula 6,7:c:1,5 using the CFSAN hqSNP pipeline. The tree was constructed with PhyML using hqSNPs identified among Illumina and ONT sequences of the strain FSL R9-0095 and 26 sequences downloaded from the NCBI. Illumina sequences of FSL R9-0095 was used as the reference genome. (a) The tree includes both the Illumina and the ONT sequencing data of strain FSL R9-0095. Clades of Typhisuis, Paratyphi C, Choleraesuis, Choleraesuis var. Kunzendorf, Choleraesuis var. Decatur are annotated separatel. (b) The data of strain FSL R9-0095 replaced with SNPs from two hrs ONT sequencing data using sequencing workflow 2. (c) The tree only includes Illumina sequencing data of strain FSL R9-0095, which were used as a benchmark phylogenetic tree. Only high support in the analysis (bootstrap >= 90%) are labeled in this tree. The tree is midpoint rooted and the scale axis is provided below the tree.

**Figure 3.**
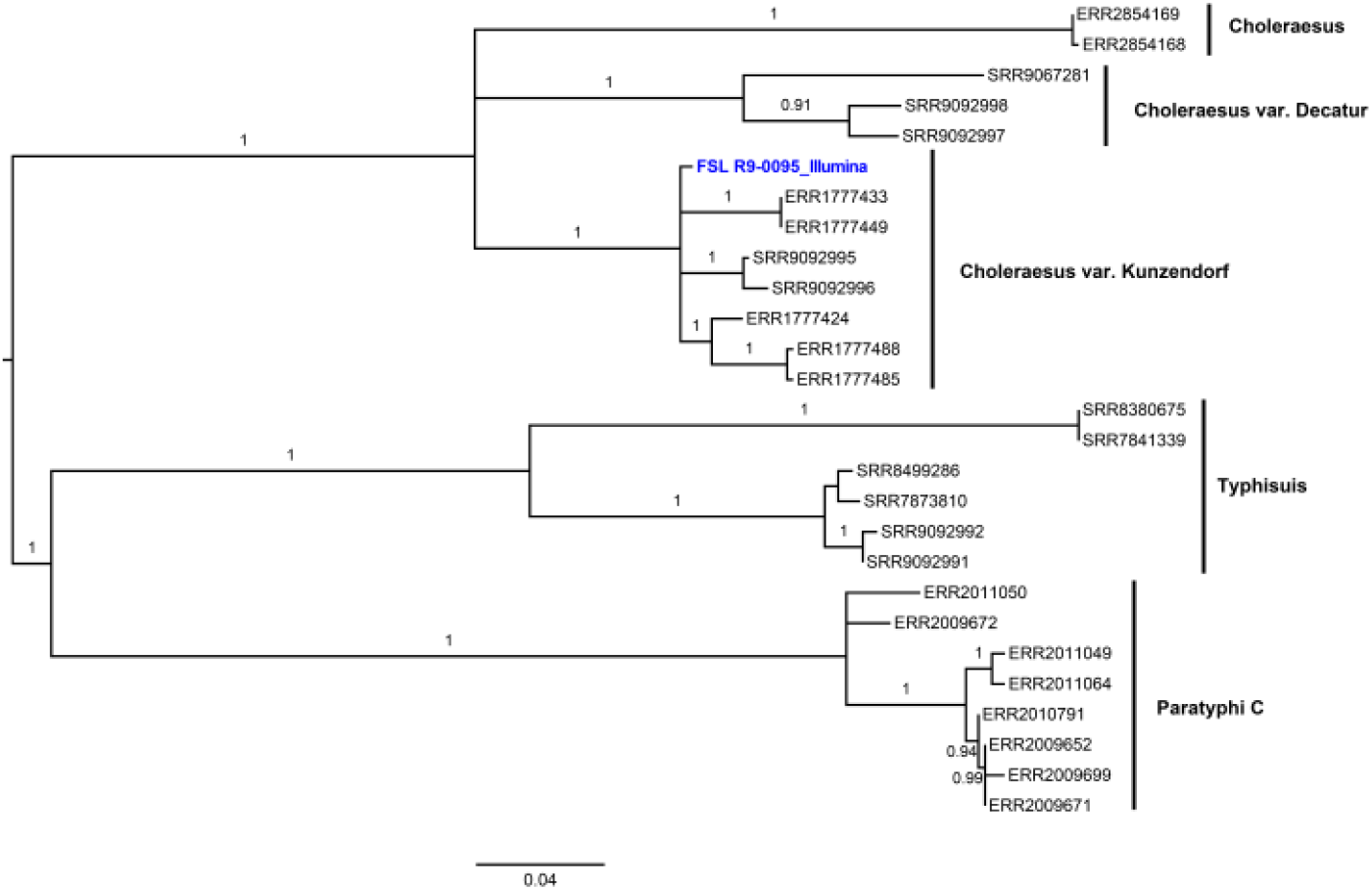

Longo et al. (2019) showed that WGS followed by Short Read Sequence Typing for Bacterial Pathogens (SRST2) for MLST (Inouye et al., 2014) in *Salmonella* TypeFinder (https://cge.cbs.dtu.dk/services/SalmonellaTypeFinder/) could correctly identify all the analyzed *S*. Choleraesuis var. Kunzendorf isolates. However, *Salmonella* TypeFinder cannot currently accept raw ONT reads as inputs and few tools are available to identify Cholerasuis var. Kunzendorf using ONT sequences. In this study, phylogenetic analysis through kSNP or CFSAN hqSNP analysis using ONT sequences successfully identified the isolate FSL R9-0095 to the variant level, which suggests the feasibility of applying ONT sequencing for differentiation of the serotypes or serotype variants sharing the same formula 6,7:c:1,5. The phylogenetic result from this study was in accordance with the result from Nair et al. (2020) using Illumina sequences.

### 3.4. Differentiation of serotype variants that differ by minor antigens

#### 3.4.1 SNP analysis

Similar results were obtained from both kSNP and CFSAN hqSNP trees generated using either the ONT or the Illumina-sequenced genomes of FSL R8-3858. All the ONT sequences were clustered with their corresponding Illumina sequences (Suplementary Figure 2 and Supplementary Figure 3a). Serotype Orion and its variants Orion var. 15^+^ and Orion var. 15^+^, 34^+^ were indistinguishable in both SNP trees (Supplementary Figure 2 and Supplementary Figure 3), indicating that phylogenetic clustering is not able to differentiate the two Orion variants from each other and Orion.

#### 3.4.2. Detection of prophage

Since the Orion variants are caused by phage conversion and phylogenetic analysis could not differentiate them, *in silico* prophage detection was used for their differention. Prophage prediction using Illumina data correctly identified Orion, Orion var. 15^+^, or Orion var. 15^+^, 34^+^ in 27 of 42 isolates (Supplementary Table 1). Four sequences were untypable or questionable since □_15_ and/or □_34_ could not be identified (Supplementary Table 1). Moreover, 11 isolates had questionable prohage prediciton that was inconsistent with the original phage typing records (Supplementary Table 1). A 25.6 Kbp region was detected with Illumina data as phage □_15_ with a score of 20 (marked as incomplete), and a 36.8 Kbp region was detected as phage □_34_ with a score of 80 (marked as questionable). Sub-optimal assembly and identification of the two phages was likely due to frequent occurrence of sequencing gaps in the prophage regions caused by short Illumina reads.

Twenty-seven of the 42 Illumina sequences tested were identified to be in accordance with their original record through PHASTER (supplymentary Table 1). By contrast, both prophages for genomes assembled from ONT data were detected with higher scores (100 – 150 for □_15_ and 80 – 90 for □_34_), except the genome assembled from 48 hr sequences using workflow 4 (Table 3). These results showed that the much longer ONT reads (compared with Illumina reads) allowed the detection of more complete prophages for definitive differentiation of Orion variants.

**Table 3.** Result of predicted prophage for strain FSL R8-3858.

| Sequencing technology | Sequencing workflow No. | Flowcell | Library preparation kit | Basecalling model | Sequencing time | Prophage □<sub>15</sub> |  |  | Prophage □<sub>34</sub> |  |  |
| --- | --- | --- | --- | --- | --- | --- | --- | --- | --- | --- | --- |
|  |  |  |  |  |  | Length (kb) | Completeness | Score | Length (bp) | Completeness | Score |
| Illumina |  |  |  |  |  | 25.6 | Incomplete | 20 | 36.8 | Questionable | 80 |
| ONT | 1 | R9 | Rapid | HAC | 2hr | 45.9 | Intact | 130 | 63.7 | Questionable | 90 |
|  |  |  |  |  | 48hr | 45.9 | Intact | 120 | 47.1 | Questionable | 80 |
|  | 2 | R9 | Rapid | Base modified | 2hr | 44.3 | Intact | 100 | 43.1 | Questionable | 90 |
|  |  |  |  |  | 48hr | 46 | Intact | 100 | 43.7 | Questionable | 80 |
|  | 3 | R9 | PCR kit | HAC | 2hr | 44.3 | Intact | 100 | 44.1 | Questionable | 90 |
|  |  |  |  |  | 72hr | 44.3 | Intact | 100 | 45.1 | Questionable | 90 |
|  | 4 | R9 | 1D2 | HAC | 2hr | 45.7 | Intact | 150 | 43.7 | Questionable | 90 |
|  |  |  |  |  | 48hr | 45.6 | Intact | 150 | 33.2 | Incomplete | 30 |
|  | 5 | R10 | Rapid | HAC | 2hr | 46 | Intact | 110 | 60.1 | Questionable | 90 |
|  |  |  |  |  | 48hr | 44.3 | Intact | 130 | 47.1 | Questionable | 90 |

**Note:** When the score was above 90, the completeness was recorded as intact; when the score was
between 70 and 90, the completeness was recorded as questionable; when the score was below 70, the
completeness was recorded as incomplete. More information about the result interpretation for scoring
the prophage region can be found on PASTER API (http://phaster.ca).

## 5. Conclusion

In this study we demonstrated that long-read nanopore sequencing technology can be used as a subtyping tool to differentiate *Salmonella* serotypes or serotype variants which share the same or related antigenic formulae. The matching identity between ONT and Illumina sequences was improved to 99.98 – 99.99% using the workflow including the bioinformatics pipeline. SNP-based phylogenetic analysis successfully identified serotypes and serotype variants including Choleraesuis, Choleraesuis var. Kunzendorf, Choleraesuis var. Decatur, Paratyphi C, Typhisuis by using ONT sequences. ONT sequencing can also successfully differentiate variants of Orion, Orion var. 15^+^ and Orion var. 15^+^, 34^+^ by addition of PHASTER prediction.

## Acknowledgement

The authors would like to thank Dr. Martin Weidmann at Cornell University for providing strains used in this study. The authors would also like to thank Oxford Nanopore Technologies (Thomas Bray, Lin Jin, Xinran Yu, Xiang Chen, Iain MacLaren, Richard Compton, Stephen Rudd, David Dai) for supporting the establishment of ONT capability at the Mars Global Food Safety Center.

