## Supplementary Tables and Figures for "Evaluation of Nanopore sequencing technology to differentiate *Salmonella* serotypes and serotype variants with the same or closely related antigenic formulae"

**Supplenmentary Table 1 Different sequencing workflows used in this study.**

| **Sequencing workflow No.** | **Library construction kit** | | **Flowcell** | | **Basecalling model** | | |
| --- | --- | --- | --- | --- | --- | --- | --- |
|  | **Name** | **Version** | **Type** | **Version** | **Basecaller** | **Version** | **Model** |
| 1 | Rapid | SQK-RAD004 | R9 | 9.4.1/FLO-MIN106 | Guppy | 3.2.6 | High accuracy (HAC) |
| 2 | Rapid | SQK-RAD004 | R9 | 9.4.1/FLO-MIN106 | Guppy | 3.2.6 | Base modified |
| 3 | PCR | SQK-PSK004 | R9 | 9.4.1/FLO-MIN106 | Guppy | 3.2.6 | High accuracy (HAC) |
| 4 | 1D2 | SQK-LSK308 | R9 | 9.5/FLO-MIN107 | Guppy | 3.2.6 | High accuracy (HAC) |
| 5 | Rapid | SQK-RAD004 | R10 | 10.0/FLO-MIN110 | Guppy | 3.2.6 | High accuracy (HAC) |

**Supplementary Table 2. Sequences from Genebank used in phylogenetic analysis. (in a separate excel file)**

**Figure legends**

**Supplementary Figure 1. Data analysis pipelines for genomic assembling and correction.** Pipeline used Wtdbg2 as the assembly tool, plus 2 rounds of correction by Racon and 1 round of Medaka.

**Supplementary Figure 2.** **Maximum parsimony tree of the serotype formulae 3,{10}{15}{15,34}:y:1,5 based on k-mer-based SNP analysis.** The tree was built using kSNP3 with the core SNPs identified in both the Illumina and ONT sequences of the isolate FSL R8-3858 and 27 sequences downloaded from the NCBI. Only high support in the analysis (bootstrap >= 90%) are labeled in this tree. The tree is midpoint rooted and the scale axis is provided below the tree.

**Supplementary Figure 3. Maximum likelihood phylogenetic tree of the serotype formulae 3,{10}{15}{15,34}:y:1,5 using the CFSAN hqSNP pipeline.** The tree was constructed with PhyML using hqSNPs identified in both the Illumina and ONT sequences of strain FSL R8-3858 and 27 sequences downloaded from the NCBI. Illumina sequences of FSL R8-3858 were used as the reference genome. (a) The tree included both the Illumina and the ONT data for strain FSL R8-3858. (b) The data for strain FSL R8-3858 were replaced with SNPs from two hrs ONT sequencing using workflow 2. (c) The tree only includes Illumina data for strain FSL R8-3858 which were used as a benchmark phylogenetic tree. Only high support in the analysis (bootstrap >= 90%) are labeled in this tree. The tree is midpoint rooted and the scale axis is provided below the tree.

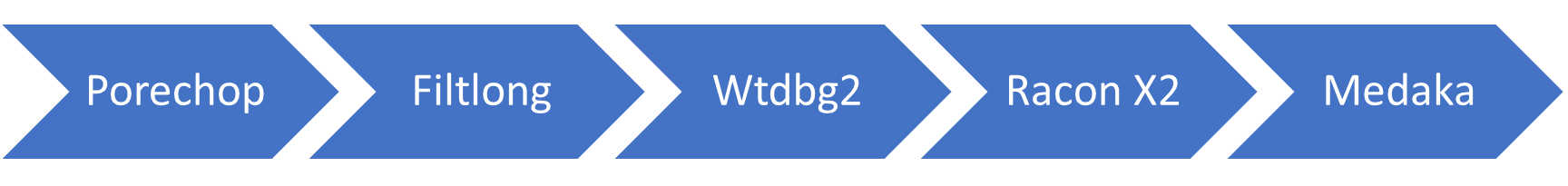

**Supplementary Figure 1**

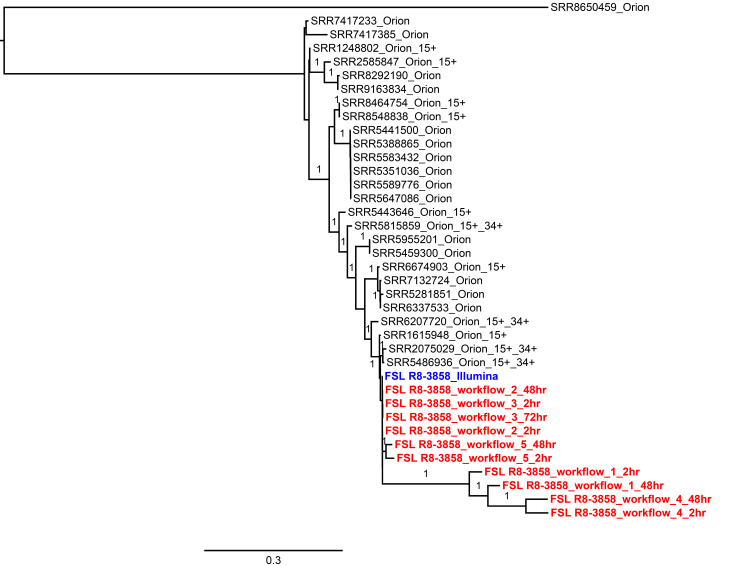

**Supplementary Figure 2**

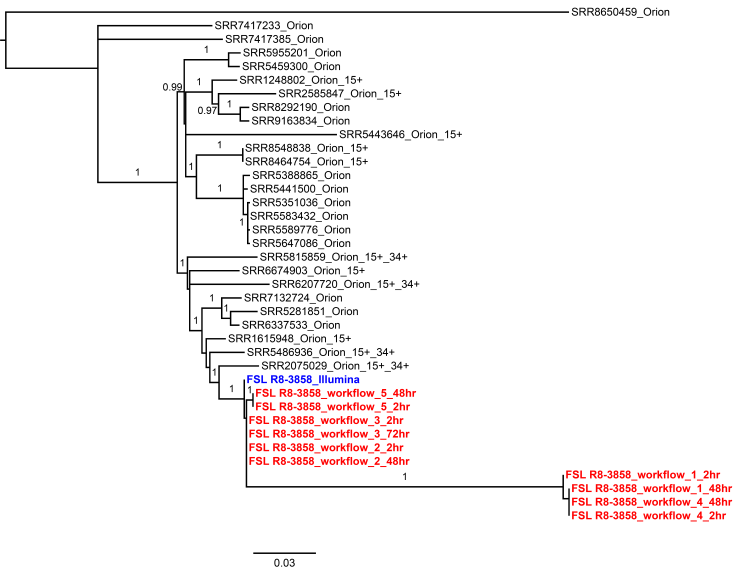

**Supplementary Figure 3a**

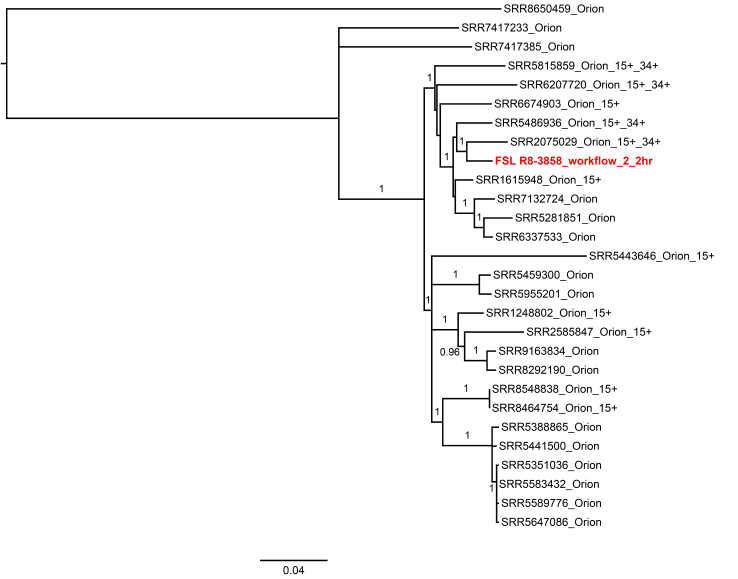

**Supplementary Figure 3b**

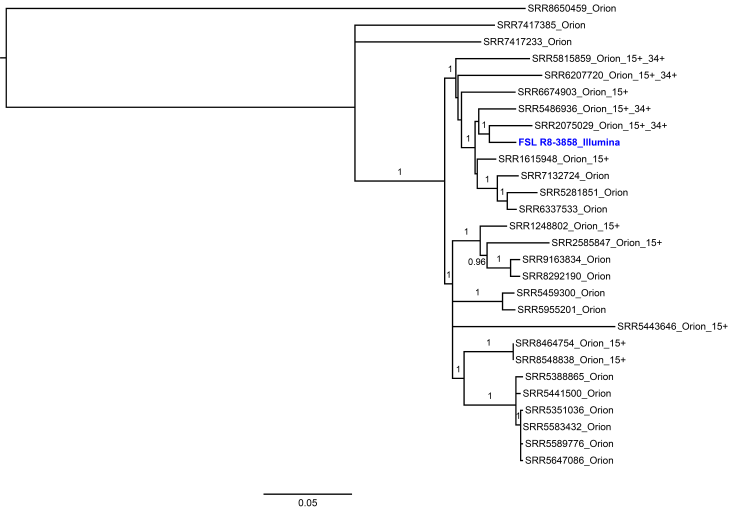

**Supplementary Figure 3c**
