## Supplementary Table 2 for "Evaluation of Nanopore sequencing technology to differentiate *Salmonella* serotypes and serotype variants with the same or closely related antigenic formulae"

**Supplementary Table 2.** Sequences from GeneBank used in phylogenetic analysis.

|  |  |  | **SISTR** | **PHASTER** | | |  |
| --- | --- | --- | --- | --- | --- | --- | --- |
| **SRA#** | **NCBI serovar name** | **SeqSero2** | **serovar** | **Ɛ15** | **Ɛ34** | **Note** | **Included in the SNP tree** |
| SRR9092995 | Choleraesuis var. Kunzendorf | Paratyphi C or Choleraesuis or Typhisuis | Choleraesuis | - | - | - | Yes |
| SRR9092996 | Choleraesuis var. Kunzendorf | Paratyphi C or Choleraesuis or Typhisuis | Choleraesuis | - | - | - | Yes |
| ERR1777488 | Choleraesuis var. Kunzendorf | Paratyphi C or Choleraesuis or Typhisuis | Choleraesuis | - | - | - | Yes |
| ERR1777424 | Choleraesuis var. Kunzendorf | Paratyphi C or Choleraesuis or Typhisuis | Choleraesuis | - | - | - | Yes |
| ERR1777485 | Choleraesuis var. Kunzendorf | Paratyphi C or Choleraesuis or Typhisuis | Choleraesuis | - | - | - | Yes |
| ERR1777433 | Choleraesuis var. Kunzendorf | Paratyphi C or Choleraesuis or Typhisuis | Choleraesuis | - | - | - | Yes |
| ERR1777449 | Choleraesuis var. Kunzendorf | Paratyphi C or Choleraesuis or Typhisuis | Choleraesuis | - | - | - | Yes |
| SRR9092997 | Choleraesuis var. Decatur | Paratyphi C or Choleraesuis or Typhisuis | Choleraesuis | - | - | - | Yes |
| SRR9092998 | Choleraesuis var. Decatur | Paratyphi C or Choleraesuis or Typhisuis | Choleraesuis | - | - | - | Yes |
| SRR9067281 | Choleraesuis var. Decatur | Paratyphi C or Choleraesuis or Typhisuis | Choleraesuis | - | - | - | Yes |
| ERR2009699 | Paratyphi C | Paratyphi C or Choleraesuis or Typhisuis | Paratyphi C | - | - | - | Yes |
| ERR2011049 | Paratyphi C | Paratyphi C or Choleraesuis or Typhisuis | Paratyphi C | - | - | - | Yes |
| ERR2011050 | Paratyphi C | Paratyphi C or Choleraesuis or Typhisuis | Paratyphi C | - | - | - | Yes |
| ERR2011064 | Paratyphi C | Paratyphi C or Choleraesuis or Typhisuis | Paratyphi C | - | - | - | Yes |
| ERR2010791 | Paratyphi C | Paratyphi C or Choleraesuis or Typhisuis | Paratyphi C | - | - | - | Yes |
| ERR2009672 | Paratyphi C | Paratyphi C or Choleraesuis or Typhisuis | Paratyphi C | - | - | - | Yes |
| ERR2009671 | Paratyphi C | Paratyphi C or Choleraesuis or Typhisuis | Paratyphi C | - | - | - | Yes |
| ERR2009652 | Paratyphi C | Paratyphi C or Choleraesuis or Typhisuis | Paratyphi C | - | - | - | Yes |
| ERR2854168 | Choleraesuis | Paratyphi C or Choleraesuis or Typhisuis | Choleraesuis | - | - | - | Yes |
| ERR2854169 | Choleraesuis | Paratyphi C or Choleraesuis or Typhisuis | Choleraesuis | - | - | - | Yes |
| SRR7841339 | Typhisuis | Paratyphi C or Choleraesuis or Typhisuis | Paratyphi C | - | - | - | Yes |
| SRR7873810 | Typhisuis | Paratyphi C or Choleraesuis or Typhisuis | Paratyphi C | - | - | - | Yes |
| SRR8380675 | Typhisuis | Paratyphi C or Choleraesuis or Typhisuis | Paratyphi C | - | - | - | Yes |
| SRR9092991 | Typhisuis | Paratyphi C or Choleraesuis or Typhisuis | Paratyphi C | - | - | - | Yes |
| SRR9092992 | Typhisuis | Paratyphi C or Choleraesuis or Typhisuis | Paratyphi C | - | - | - | Yes |
| SRR8499286 | Typhisuis | Paratyphi C or Choleraesuis or Typhisuis | Paratyphi C | - | - | - | Yes |
| SRR9163834 | Orion | Orion | Orion | NA | NA | Correctly identified | Yes |
| SRR8650459 | Orion | Orion | Orion | NA | NA | Correctly identified | Yes |
| SRR8292190 | Orion | Orion | Orion | NA | NA | Correctly identified | Yes |
| SRR7417385 | Orion | Orion | Orion | NA | NA | Correctly identified | Yes |
| SRR7417233 | Orion | Orion | Orion | NA | NA | Correctly identified | Yes |
| SRR7132724 | Orion | Orion | Orion | NA | NA | Correctly identified | Yes |
| SRR6337533 | Orion | Orion | Orion | NA | NA | Correctly identified | Yes |
| SRR5955201 | Orion | Orion | Orion | NA | NA | Correctly identified | Yes |
| SRR5647086 | Orion | Orion | Orion | NA | NA | Correctly identified | Yes |
| SRR5589776 | Orion | Orion | Orion | NA | NA | Correctly identified | Yes |
| SRR5583432 | Orion | Orion | Orion | NA | NA | Correctly identified | Yes |
| SRR5459300 | Orion | Orion | Orion | NA | NA | Correctly identified | Yes |
| SRR5441500 | Orion | Orion | Orion | NA | NA | Correctly identified | Yes |
| SRR5388865 | Orion | Orion | Orion | NA | NA | Correctly identified | Yes |
| SRR5351036 | Orion | Orion | Orion | NA | NA | Correctly identified | Yes |
| SRR5281851 | Orion | Orion | Orion | NA | NA | Correctly identified | Yes |
| SRR8464754 | Orion var. 15+ | Orion | Orion | 20 | NA | Correctly identified | Yes |
| SRR8548838 | Orion var. 15+ | Orion | Orion | 20 | NA | Correctly identified | Yes |
| SRR6674903 | Orion var. 15+ | Orion | Orion | 100 | NA | Correctly identified | Yes |
| SRR5443646 | Orion var. 15+ | Orion | Orion | 100 | NA | Correctly identified | Yes |
| SRR2585847 | Orion var. 15+ | Orion | Orion | 110 | NA | Correctly identified | Yes |
| SRR1615948 | Orion var. 15+ | Orion | Orion | 100 | NA | Correctly identified | Yes |
| SRR1248802 | Orion var. 15+ | Orion | Orion | 20 | NA | Correctly identified | Yes |
| SRR2075029 | Orion var. 15+ 34+ | Orion | Orion | 20 | 80 | Correctly identified | Yes |
| SRR6207720 | Orion var. 15+ 34+ | Orion | Orion | 20 | 98 | Correctly identified | Yes |
| SRR5815859 | Orion var. 15+ 34+ | Orion | Orion | 20 | 90 | Correctly identified | Yes |
| SRR5486936 | Orion var. 15+ 34+ | Orion | Orion | 100 | 90 | Correctly identified | Yes |
| SRR4175484 | Orion | Orion | Orion | 108 | NA | Questionable | No |
| SRR6818878 | Orion | Orion | Orion | 20 | NA | Questionable | No |
| SRR7185127 | Orion | Orion | Orion | 20 | NA | Questionable | No |
| SRR7889327 | Orion | Orion | Orion | 20 | NA | Questionable | No |
| SRR8248606 | Orion | Orion | Orion | 120 | NA | Questionable | No |
| SRR8957106 | Orion | Orion | Orion | 100 | NA | Questionable | No |
| SRR9984452 | Orion | Orion | Orion | 20 | NA | Questionable | No |
| SRR9984455 | Orion | Orion | Orion | 20 | NA | Questionable | No |
| SRR2087275 | Orion var. 15+ | Orion | Orion | 20 | 70 | Questionable | No |
| SRR2131384 | Orion var. 15+ | Orion | Orion | 20 | 70 | Questionable | No |
| SRR2728263 | Orion var. 15+ | Orion | Orion | NA | NA | Untypable or questionable | No |
| SRR3219073 | Orion var. 15+ | Orion | Orion | 20 | 70 | Questionable | No |
| SRR3057158 | Orion var. 15+ 34+ | Orion | Orion | 100 | NA | Untypable or questionable | No |
| SRR5223595 | Orion var. 15+ 34+ | Orion | Orion | 100 | NA | Untypable or questionable | No |
| SRR7946486 | Orion var. 15+ 34+ | Orion | Orion | 108 | NA | Untypable or questionable | No |
